# MRS-measured Glutamate versus GABA reflects excitatory versus inhibitory neural activities in awake mice

**DOI:** 10.1101/2020.10.25.353888

**Authors:** Yuhei Takado, Hiroyuki Takuwa, Kazuaki Sampei, Takuya Urushihata, Manami Takahashi, Masafumi Shimojo, Shoko Uchida, Nobuhiro Nitta, Sayaka Shibata, Keisuke Nagashima, Yoshihiro Ochi, Maiko Ono, Jun Maeda, Yutaka Tomita, Naruhiko Sahara, Jamie Near, Ichio Aoki, Kazuhisa Shibata, Makoto Higuchi

## Abstract

To assess if magnetic resonance spectroscopy (MRS)-measured Glutamate (Glu) and GABA reflect excitatory and inhibitory neural activities, respectively, we conducted MRS measurements along with two-photon mesoscopic imaging of calcium signals in excitatory and inhibitory neurons of living, unanesthetized mice. For monitoring stimulus-driven activations of a brain region, MRS signals and mesoscopic neural activities were measured during two consecutive sessions of 15-min prolonged sensory stimulations. In the first session, putative excitatory neuronal activities were increased, while inhibitory neuronal activities remained at the baseline level. In the second half, while excitatory neuronal activities remained elevated, inhibitory neuronal activities were significantly enhanced. We also assessed regional neurochemical and functional statuses related to spontaneous neural firing by measuring MRS signals and neuronal activities in a mouse model of Dravet syndrome under a resting condition. Mesoscopic assessments showed that activities of inhibitory neurons in the cortex were diminished relative to wild-type mice in contrast to spared activities of excitatory neurons. Consistent with these observations, the Dravet model exhibited lower concentrations of GABA than wild-type controls. Collectively, the current investigations demonstrate that the MRS-measured Glu and GABA can reflect spontaneous and stimulated activities of neurons producing and releasing these neurotransmitters in an awake condition.

## Introduction

Proton magnetic resonance spectroscopy (MRS) is a noninvasive technique that can measure the concentrations of glutamate (Glu; chief excitatory neurotransmitter) and gamma-aminobutyric acid (GABA; chief inhibitory neurotransmitter) within a region of interest in the brain. Since the first human application in the 1980s, there has been a growing interest in the use of MRS to correlate cognition and behavior with the concentrations of Glu and GABA in specific brain regions ^1-5^. Despite the ever-greater amount of findings based on MRS, one open question remains unsolved: do MRS-measured Glu and GABA in a brain region reflect activities of excitatory and inhibitory neurons, respectively, in the region? This uncertainty has been one of the main limitations to understanding the relationships between behavior and brain processing that underly MRS signals. While there have been several studies in which brain activity, as measured by non-invasive function measures such as functional MRI (fMRI) ^6^ or magnetoencephalography (MEG) ^7^, have been related to MRS neurotransmitter measures, to our knowledge, there has been no neuroscientific study directly addressed this question by monitoring neural activities at the cellular level.

There is numerous evidence which showed the significant influence of anesthesia on neural activity ^8, 9^. Despite the importance of the awake condition for investigating neural activity and brain function, to our knowledge, no MRS work in rodents has been performed in an awake condition. Thus, to satisfactorily answer the above question, it is essential to establish a link between neural activities and MRS signals in awake and behaving conditions.

This study aims to test the hypothesis that MRS-measured Glu and GABA concentrations reflect activities of excitatory and inhibitory neurons, respectively, in an awake condition. We specifically focused on two types of awake conditions: task and quiet rest. Several studies in cognitive neuroscience have measured MRS signals in association with certain tasks ^1-4, 10, 11^. On the other hand, clinical studies have measured MRS signals mainly at quiet rest ^9, 12-14^. If the hypothesis is correct, MRS-measured Glu and GABA should reflect both task-evoked and spontaneous activities of excitatory and inhibitory neurons, respectively.

To test the hypothesis, we utilized the following two cutting edged modalities: a MRS compatible awake mouse restraint device and a large-field-of-view two-photon mesoscope with subcellular resolution, which has the advantage for measuring neural activities from a large number of neurons simultaneously ^15^. In a task condition (Experiment 1), we measured time-courses of calcium (Ca) signals and MRS during a 30-min whisker stimulation, since MRS-measured Glu and GABA in brain areas were found to be changed over the course of 20-30-min task loads ^1, 16^. To examine the relationships between MRS signals and neural activities at quiet rest (Experiment 2), we used a mouse model of Dravet syndrome, which is a severe epileptic encephalopathy from de novo mutations of SCN1A ^17^. It is known that activities of inhibitory neurons are specifically reduced in these mice compared to wild-type mice ^17^, allowing us to test whether the GABA concentration, but not the Glu concentration, is specifically decreased in these model mice.

## Material and Methods

### Animals

#### C57BL/6J

All mice were housed individually in separate cages with water and food ad libitum. Mouse cages were kept at a temperature of 25°C in a 12-h light/dark cycle. Prior to two-photon imaging and MRI/MRS scanning, all mice were handled daily for 5 days (5–10 min/day) to minimize handling stress.

#### A mouse model of Dravet syndrome

Balb/c-Scn1a<+/−> mouse (Dravet mouse) was provided by RIKEN BRC through the National Bio Resource Project of the MEXT, Japan (BRC No.: RBRC06422) ^18^. For in vivo two-photon imaging, heterozygous mutant and wild-type control mice (6 – 7 months old) were generated by crossing once with the C57BL/6 background.

All animal experiments were approved by the Institutional Animal Care and Use Committee of the National Institute of Radiological Sciences (Chiba, Japan) and were performed in accordance with the institutional guidelines on humane care and use of laboratory animals approved by the Institutional Committee for Animal Experimentation.

### MRI/MRS-compatible awake mouse restraint device and whisker stimulation

To measure the levels of neurochemicals from an awake mouse by MRS, we developed an MRI/MRS compatible awake mouse restraint device to hold the skull of the mouse tightly on the holder (Fig. 1A). The fixation method of a mouse head is in principle identical to the one for two-photon microscopy ^19^. The surgery was performed two weeks before the MRI and MRS measurements, then after recovery from the surgery, mice were habituated for the mouse restraint device for five days. The homemade holder for the MRI scanner was optimized for adjusting the space in the bore of the 7T magnet with cryoprobe for mice. Whisker stimulation, by the use of air-puffs is available on this holder (Fig. 1B). To give sufficient stimulation to the mouse brain, multiple whiskers on the right side were stimulated together using the air-puff for 1 sec at a rate of once every 3 sec.

**Fig. 1.**
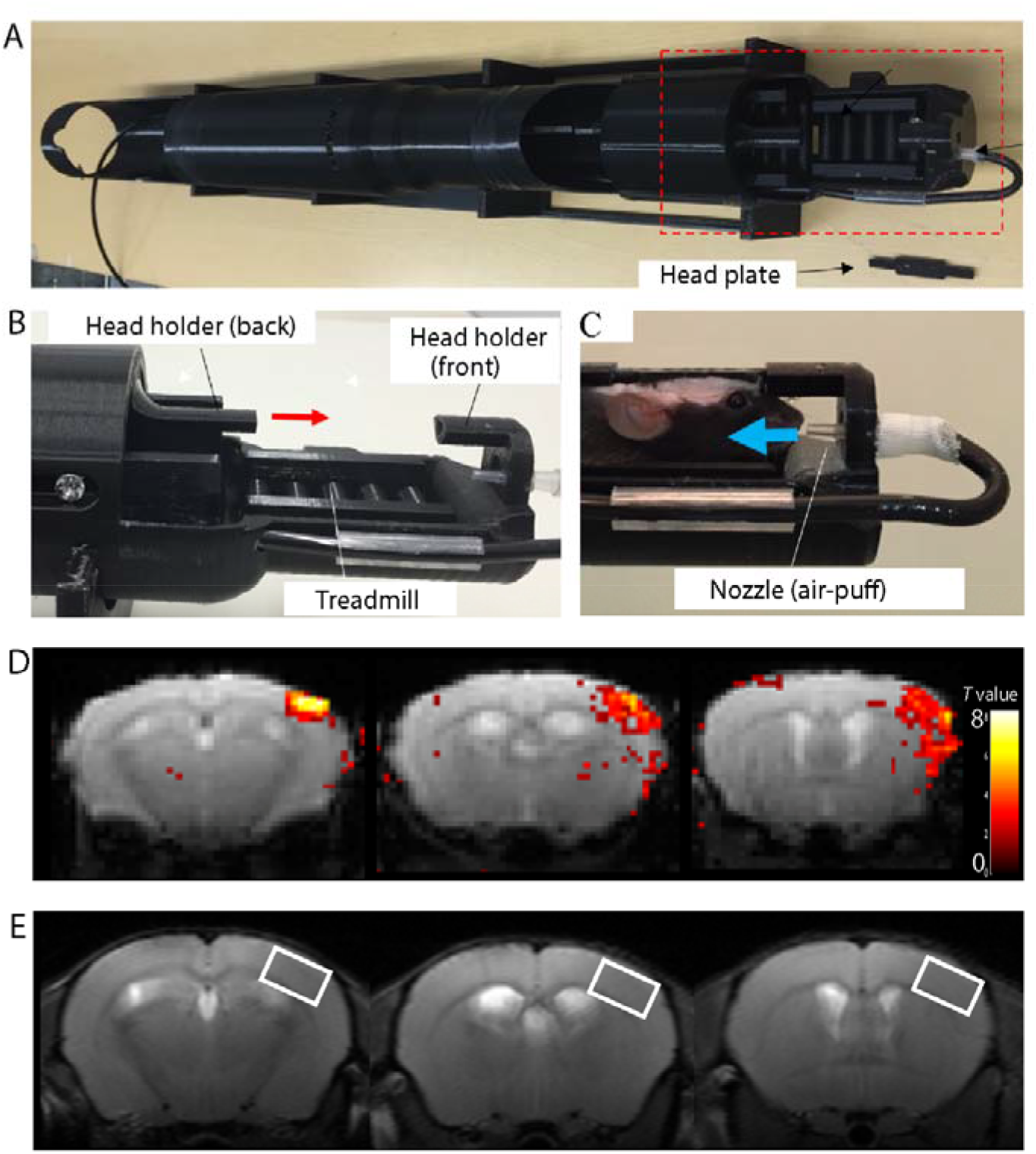
Pictorial features of the MRI/MRS-compatible awake mouse restraint device. (A) The overview of the MRI/MRS-compatible awake mouse restraint device mouse holder is shown. The region within the red-dotted rectangle is magnified in B. (B, C) Two parts (front and back) of the head plate, which is fixed on the mouse head, are inserted into the mouse holder to prevent movement of the mouse. (C) Whisker stimulation by air puff is possible for a mouse on the holder. (D) BOLD signals during whisker stimulation are shown on the barrel cortex. (E) A representative voxel placement on the barrel cortex for MRS is shown.

### MRI and MRS Measurements

#### C57BL/6J

MRI and MRS were performed on a 7.0 T MRI scanner (20 cm bore, Biospec, Avance-III system; Bruker Biospin, Ettlingen, Germany) with a 2-channel phased-array cooled surface coil for transmission/reception (cryoprobe for mouse brain, Bruker Biospin). For awake mouse experiments, mice were lightly anesthetized (1–2% isoflurane in air; 50 : 50 at 1 L/min) for < 3 min while being placed in the MRI/MRS-compatible awake mouse restraint device. After moving the mouse on the holder, anesthesia was turned off. Axial and sagittal multi-slice TurboRARE T2-weighted images (Axial: TR = 3000 ms, TE = 12 ms (effective TE = 36 ms), Rare Factor = 8, number of averages = 1, number of slices = 20, FOV = 19.2 mm × 19.2 mm, matrix = 256 × 256; Sagittal: TR = 3000 ms, TE 10 ms (effective TE = 30 ms), Rare Factor = 8, number of averages = 1, number of slices = 13, FOV = 19.2 mm × 12.8 mm, matrix = 128) were acquired. To confirm that the VOI of MRS in the barrel cortex is appropriately localized, functional MRI (fMRI) was performed under whisker stimulation at a rate of 2 Hz with 1-sec stimulation using a gradient-echo echo-planar sequence (TR = 500 ms, TE = 15 ms, BW = 250 kHz, number of averages = 2, number of slices = 13, spatial resolution = 200 μm × 200 μm × 1000 μm, FOV = 19.2 × 19.2 mm, matrix = 96 × 96). The stimulus was delivered using a block design paradigm of 100-sec rest and 20-sec activation alternately repeated five times. Magnetic resonance spectra were acquired from a VOI centered in the left barrel cortex. The size of the voxel was 1.175 mm × 2.0 mm × 3.5 mm. Localized MRS was applied using Point Resolved Spectroscopy (PRESS) with TE/TR = 11/3000 ms, 292 averages. MRS measurements were repeated three times continuously. At first, MRS scan was performed without whisker stimulation. Then, two more MRS scans (1st half and 2nd half) were performed with whisker stimulation at 0.333 Hz with 1-s pulse duration and 2-sec interval. The MRS protocol is shown in Fig. 3A. For experiments under isoflurane anesthesia, induction was performed by using 3% anesthesia, and anesthesia then was maintained with 1.5% isoflurane during the experiments. The rest of the manipulations were the same between awake mice and anesthetized mouse experiments.

#### A mouse model of Dravet syndrome

Heterozygous mutant and wild-type mice (2-6 months old) were used for MRS measurements in the awake condition. Magnetic resonance spectra were acquired from a VOI centered in the middle frontal cortex. The voxel size was 1.2 mm × 2.5 mm × 3.0 mm. Localized MRS was applied using PRESS with TE/TR = 11/4000 ms, 192 averages.

### fMRI Data analysis

fMRI images were created using a statistical parametric mapping package (SPM8; www.fil.ion.ucl.ac.uk/spm). Image processing steps, including (1) realigning and reslicing all EPI images within a time-series, based on their averaged image, to minimize movement artifacts, and (2) smoothing the images by using a Gaussian kernel with full width at half maximum (FWHM) of 0.5 mm, were applied to the raw data. Statistical analysis was conducted using general linear modeling with a hemodynamic response function. To control the probability of false positive clusters below 0.05, we determined the significant BOLD response areas with an individual voxel threshold of P < 0.05. This analysis was performed after the fMRI experiment.

### MRS data analysis

Acquired spectra were analyzed using LCModel (Stephen Provencher Inc, Oakville, Ontario, Canada). The unsuppressed water signal measured from the same VOI was used as an internal reference for the absolute quantification of metabolites. Data were fitted to a linear combination of 21 metabolites in a simulated basis set containing: alanine, aspartate (Asp), phosphorylcholine (PCh), creatine (Cre), phosphocreatine (PCr), GABA, glutamine (Gln), Glu, glutathione (GSH), glycine (Gly), myo-inositol (mI), lactate (Lac), N-acetylaspartate (NAA), Scyllo-inositol (Scyllo), taurine (Taur), glucose (Glc), N-acetylaspartylglutamate (NAAG), glycerophosphorylcholine (GPC), phosphorylethanolamine (PE), serine (Ser) and β-hydroxybutyrate (bHB). The average Cramér-Rao lower bounds (CRLB) values of the following 10 metabolites were < 15%: Asp, GABA, Gln, Glu, GSH, mI, Taur, total choline (tCho; the sum of GPC and PCh), total NAA (tNAA; the sum of NAA and NAAG) and total Cre (tCre; the sum of Cre and PCre). Since we focused on excitatory and inhibitory neuronal activities, Glu and GABA were utilized for the further analysis.

### Plasmid construction and AAV preparation

cDNA encoding GCaMP6s, an ultra-sensitive genetically encoded fluorescent calcium indicator ^20^, was amplified by polymerase chain reaction from pGP-CMV-GCaMP6s plasmid (a gift from Douglas Kim, Addgene plasmid # 40753) and inserted into the multi-cloning site of Adeno Associated Virus (AAV) transfer plasmid with rat synapsin promoter (pAAV-Syn), woodchuck posttranscriptional regulatory element (WPRE), and polyA signal flanked with ITRs. To generate AAV plasmid drive mCherry expression in inhibitory neurons, the promoter was substituted by mDlx enhancer sequence from pAAV-mDlx-GFP-Fishell-1 (a gift from Gordon Fishell, Addgene plasmid # 83900) and cDNA encoding mCherry was further inserted into multicloning site ^21^. For large scale preparation of recombinant AAV, each AAV transfer plasmid and AAV serotype DJ packaging plasmids (pHelper and pRC-DJ) were introduced into HEK293T cells with polyethyleneimine transfection. 48h after transfection, cells were harvested, lysed, and AAV particles were purified with HiTrap heparin column (GE Healthcare) as described previously ^22^.

### Wide-field two-photon imaging in mice

AAV vector driving GCaMP6s gene (AAV-Synapsin) and DLX (AAV-DLX) were co-injected into the barrel cortex two months before the experiment with two-photon imaging. For surgery, the animals were anesthetized with a mixture of air, oxygen, and isoflurane (3–4% for induction and 2% for surgery) via a facemask, and a cranial window (3-4 mm in diameter) was attached over the left somatosensory cortex, centered at 1.8mm caudal and 2.5mm lateral from the bregma, according to the ‘Seylaz-Tomita method’ ^23^. All two-photon imaging experiments were performed four weeks after the cranial window surgery. The method for preparing the chronic cranial window was previously reported elsewhere ^19, 24^. The awake animal was placed on a custom-made apparatus, and real-time imaging was conducted by wild-field two-photon laser scanning microscopy (Multiphoton Mesoscope, THORLABS, NJ, USA). In this study, two laser oscillators were used to efficiently separate red and green fluorescence (Supplementary Fig.4). One is a commercially available laser oscillator (Mai Tai, Spectra-Physics, CA, USA) and the other, a Yb-doped fiber laser system, was used as a pump laser for the two-photon microscope. The latter laser system is home-made and consisted of a mode-locked fiber oscillator, multi-stage fiber amplifiers and a pulse compressor. We used a nonlinear amplification method ^25, 26, 27^ in the amplifiers. The laser had a pulse repetition frequency of 73 MHz, a central wavelength of 1036 nm, a pulse length (full width at half maximum) of 130 fs and an output power of 5 W. Excitation wavelength of 900 nm (commercial laser oscillator) and 1036 nm (home-made laser system) were used for simultaneous measurements of GCaMP and DLX, respectively. An emission signal was separated by a beam splitter (560/10 nm) and simultaneously detected with a band-pass filter for GCaMP (525/50 nm) and mCherry (610/75 nm). The visual field size of the image was 500 μm × 500 μm and 3000 μm × 3000 μm, and in-plane pixel size was 1 µm. Temporal resolution was 1-4 Hz for 60 sec (60 - 240 frames/trial) depending on the size of the field of view. Images were acquired at a depth of about 150 μm from the brain surface.

In the case of Supplementary Fig. 1 (500 μm × 500 μm image), ten trials were successively performed with an inter-trial interval of 60 sec. After 30 sec from the start of the scan, activation of neurons was induced by sensory stimulation with an air puff for 10 sec. In case of Fig. 4 (3000 μm × 3000 μm image), 60-sec images were made 15 times (total 30 min), once per min. Immediately after the start of the scan, air puff stimulation was given continuously for 30 min, once every 3 sec. The patterns of whisker stimulation were the same condition as for the MRS experiments.

Two-photon imaging was performed in awake mice. The experimental protocol for measurements by two-photon imaging was reported previously ^8, 24^. Briefly, the handmade fixing device on the animal’s head was screwed to a custom-made stereotactic apparatus. The animal was then placed on a styrofoam ball that was floating using a stream of air. This allowed the animal to exercise freely on the ball while its head was fixed to the apparatus.

### Image analysis of two-photon microscopy

Image analysis of two-photon calcium imaging was performed using Matlab (MathWorks, MA, USA). For analysis of sensory stimulation experiments, data on the timing when the mouse moved were first removed. Next, motion correction was performed using NoRMCorre. The images for all time points were divided by the first image, and the images of signal change rate were obtained. Region of interests (ROIs) were manually drawn on the image obtained by maximum intensity projection (MIP) processing for images of the sensory stimulation period. The time-dependent change in average luminance value was then obtained for each ROI.

### Statistical analyses

Results are presented as mean ± standard deviation. Normality testing was performed, all data passed normality (Shapiro-Wilk test), and then parametric statistical analysis was performed. Data were analyzed in OriginPro 2019 (OriginLab, Northampton, MA, USA) with two-sample, paired t-test or one-way repeated-measures ANOVA with Bonferroni post-hoc test, when appropriate. P < 0.05 was considered significant.

## Results

### Differential neural activities at awake and anesthetic conditions

We first confirmed this assumption in our study: neural activities in the awake condition are different from those in the anesthetic condition. 1-Hz whisker stimulation, administered via air-puffs, was performed for 10 sec on a mouse while its skull was tightly fixed by a homemade awake mouse restraint device (Fig. 1A-C). Two-photon Ca imaging of neurons in the barrel cortex was performed on the head-fixed mice during awake condition and under anesthesia (isoflurane)^8^. While no significant increase of Ca signal intensities in neurons was found in the anesthetic condition (Supplementary Fig. 1A, B left), we found an increase in Ca signal intensities in the awake condition in response to whisker stimulation (Supplementary Fig. 1A, B right). These differences clearly indicate that neural activities are different between the awake and anesthetic conditions.

Secondly, we identified the area of the barrel cortex that responds to whisker stimulation using functional magnetic resonance imaging (fMRI)^28^. As shown in Fig. 1D, the region in the somatosensory cortex was activated by whisker stimulation. Based on this data, we localized a voxel of interest (VOI) that was used for the following MRS measurements (Fig. 1E).

### Activities of excitatory and inhibitory neurons changed in accordance with changes in MRS-measured Glu and GABA concentrations, respectively, during prolonged whisker stimulation

To test whether MRS-measured Glu and GABA reflect excitatory and inhibitory neural activities, respectively, we applied the identical whisker stimulation protocol to both two-photon imaging and MRS experiments. To provoke changes of excitatory and inhibitory neural activities in the barrel cortex (Fig. 2A), we applied whisker stimulation (1-sec air-puff at 0.333 Hz frequency) for 30-min. The time duration was designed based on previous reports in which MRS-measured Glu and GABA in the cortex could be differentially altered over the course of 20-30-min task loads ^1, 16^. Before the stimulation, 10- to 15-min baseline measurements were performed in both two-photon imaging and MRS measurements. Data obtained from this pre-stimulation period served as a baseline.

**Fig. 2.**
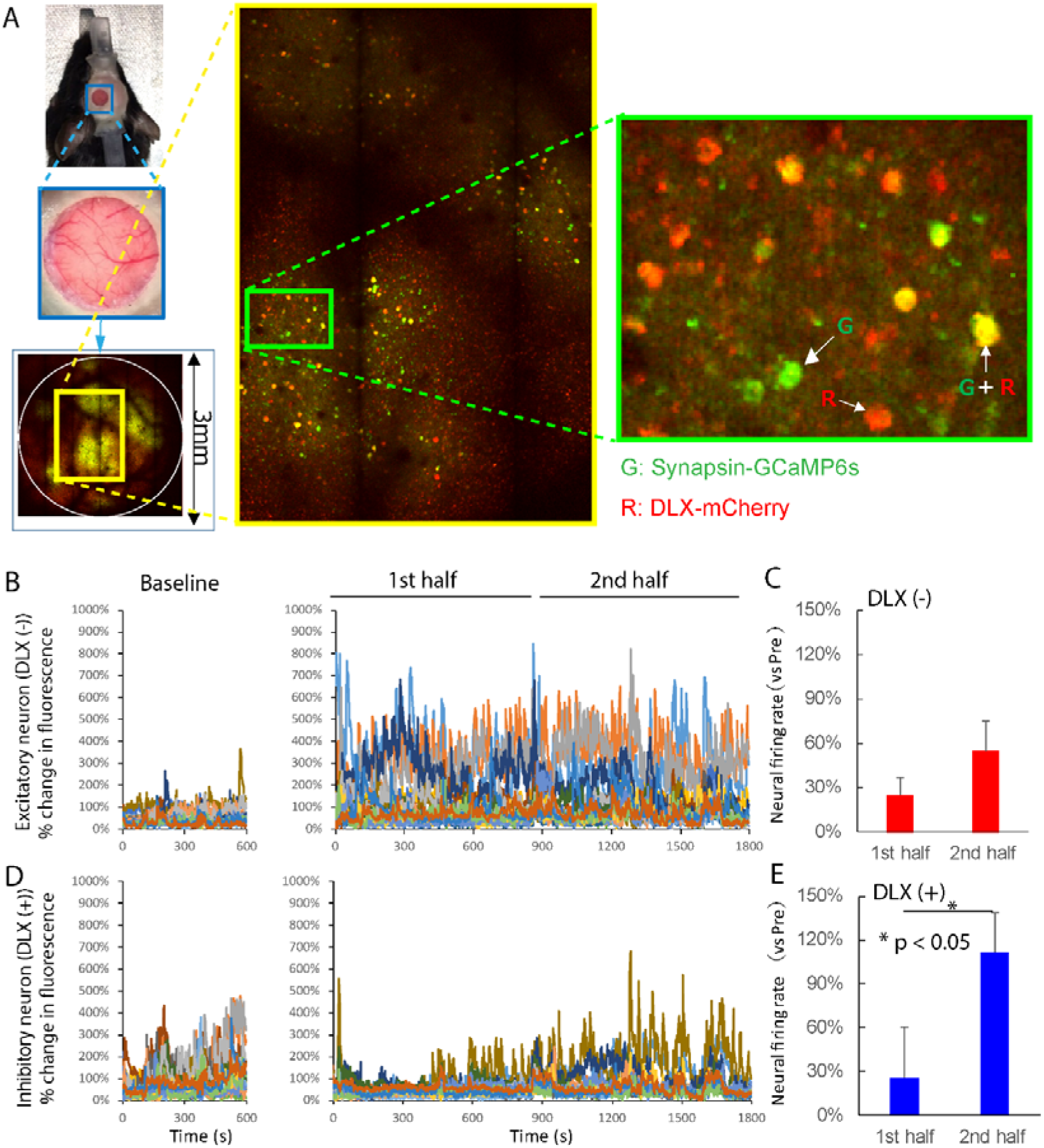
Neural activities measured by two-photon laser microscopy at the awake condition. (A) A large field-of-view of the two-photon mesoscope is shown. Calcium indicator GCaMP6s driven by Synapsin 1 promoter were utilized to express specifically in neuron cells. To generate AAV plasmid drive mCherry expression in inhibitory neurons, the promoter was substituted by mDlx enhancer sequence. By overlapping both colors, inhibitory neural activity could be deduced by yellow colored cells. After 10 min of two-photon imaging without stimulation, 30-min scans were conducted; the 1st half for 15 min and the 2nd half for 15 min by the same protocol as for the MRS measurements. At the 1st half session of whisker stimulation, Ca signals of putative excitatory neurons became evident (B, C) and inhibitory neurons remained small (D, E). At the 2nd half session of whisker stimulation, Ca signals of putative excitatory neurons remained evident (B, C) and inhibitory neurons became significantly higher than in the 1st half session (paired t-test, n = 4 mice, t = −3.3536, p = 0.044) (D, E). Error bars indicate standard deviation.

During the first half of whisker stimulation (15 min), activities of putative excitatory neurons were increased from the baseline (Fig. 2B, C). The activities stayed at an increased level in the second half, resulting in insignificant difference between the first and second half sessions (paired t-test, n = 4 mice, t = −1.971, p = 0.143). On the other hand, the activities of inhibitory neurons became increased from the baseline only during the second half (paired t-test, n = 4 mice, t = −3.354, p = 0.044) (Fig.2D, E). These results indicate that the prolonged whisker stimulation protocol successfully led to changes in activities of excitatory and inhibitory neurons at the different time points.

Using the same stimulation protocol, we performed MRS to confirm the link between changes in excitatory and inhibitory neural activities and changes in the concentrations of MRS-measured Glu and GABA, respectively. Three 15-min MRS sessions were conducted consecutively (Fig.3A) ⍰ the baseline acquisition without stimulation, and the first half and second half acquisitions with whisker stimulation. Representative spectra (Fig.3B) and GABA fitted spectra in the awake condition (Fig. 3C) were shown. All of the fitted spectra by LCModel^29^ are presented in Supplementary Fig.2.

In the first half of the stimulation (15 min), a significant increase of Glu levels was observed (one-way repeated-measures ANOVA with Bonferroni post-hoc test (rm-ANOVA), n = 8, t = 4.288, p = 0.002) without significant alteration of the GABA levels (rm-ANOVA, n = 8, t = 1.505, p = 0.155). In the second half, the GABA levels were increased significantly compared to baseline (rm-ANOVA, n = 8, t = 3.589, p = 0.003), while the Glu levels did not increase further (Fig. 3D, E). These results indicate that changes of MRS-measured Glu and GABA concentrations were consistent with changes of excitatory and inhibitory neural activities, respectively, in the awake condition. In contrast to those findings, Glu and GABA levels in the anesthetic condition did not increase by whisker stimulation, and in fact they rather decreased in the second half (Glu; rm-ANOVA, n = 8, t = 3.055, p = 0.030, GABA; rm-ANOVA, n = 8, t = 2.737, p = 0.018), which may be attributable to the lack of brain fuel due to anesthesia (Supplementary Fig.3).

**Fig. 3.**
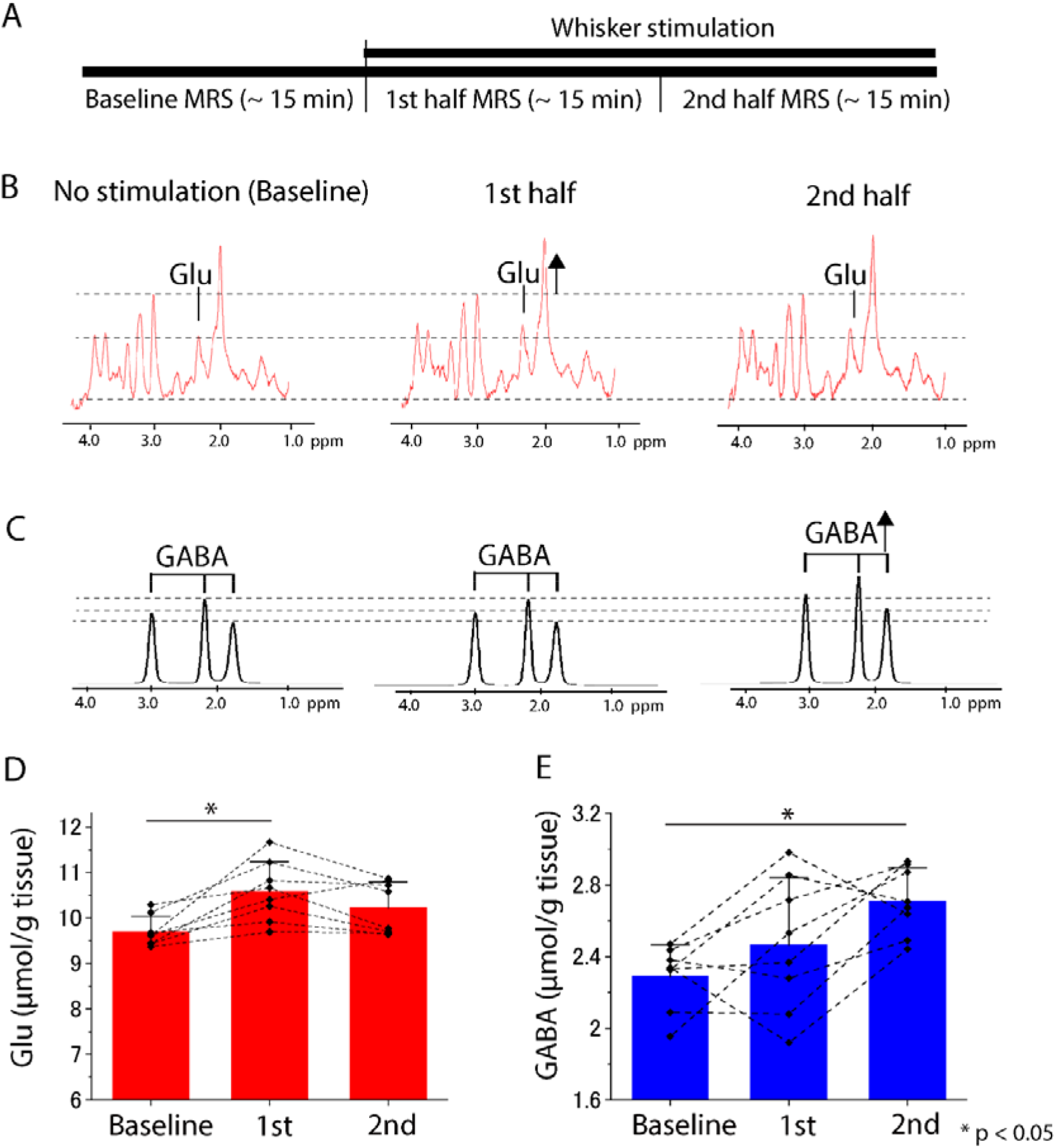
Alterations of Glu and GABA levels during whisker stimulation measured by MRS. (A) Three MRS acquisitions (one baseline measurement and two measurements with whisker stimulation) were performed in the awake condition. The MRS protocol is shown in (A). (B,C) Representative spectra in the awake condition are shown. GABA signals after LCModel fitting are also shown at the lower middle. Significant alterations of Glu and GABA were found during the awake condition. In the first half of the stimulation (15 min), a significant increase of Glu levels was observed (one-way repeated-measures ANOVA with Bonferroni post-hoc test (rm-ANOVA), n = 8, t = 4.288, p = 0.002) without significant alteration of the GABA levels (rm-ANOVA, n = 8, t = 1.505, p = 0.155). In the second half, the GABA levels were increased significantly compared to baseline (rm-ANOVA, n = 8, t = 3.589, p = 0.003) whereas the Glu levels decreased slightly (D, E). Quantitative alterations of Glu and GABA at the awake condition are shown. Error bars indicate standard deviation.

### Weaker activities of inhibitory neurons are associated with lower concentrations of MRS-measured GABA during quiet rest

Next, we tested whether MRS-measured Glu and GABA during quiet rests reflect spontaneous activities of excitatory and inhibitory neurons, respectively. We employed an epilepsy mouse model of Dravet syndrome, which is known to have dysfunction of inhibitory neurons due to a gene mutation of SCN1A^17^. We found no significant changes in spontaneous activities of excitatory neurons measured by two-photon microscopy (two-sample t-test, n = 219-356 cells, t = 0.9315, p = 0.3519; Fig. 4B, C) or MRS-measured Glu levels (two-sample t-test, n = 8, t = 0.066, p = 0.948; Fig. 4F) between the model and wild-type mice. On the other hand, we found that spontaneous activities of inhibitory neurons in the model mice were significantly lower than those in wild-type mice (Two-sample t-test, n = 41-61 cells, t = −4.168, p = 6.628E-5; Fig. 4D, E). Consistent with these decreased activities, the model mice also showed lower GABA levels in gray matter than wild-type mice at the resting state (two-sample t-test, n = 7, t = 3.010, p = 0.008; Fig. 4C). These results indicate that MRS-measured Glu and GABA reflect excitatory and inhibitory neural activities, respectively, at quiet rest condition.

**Fig. 4.**
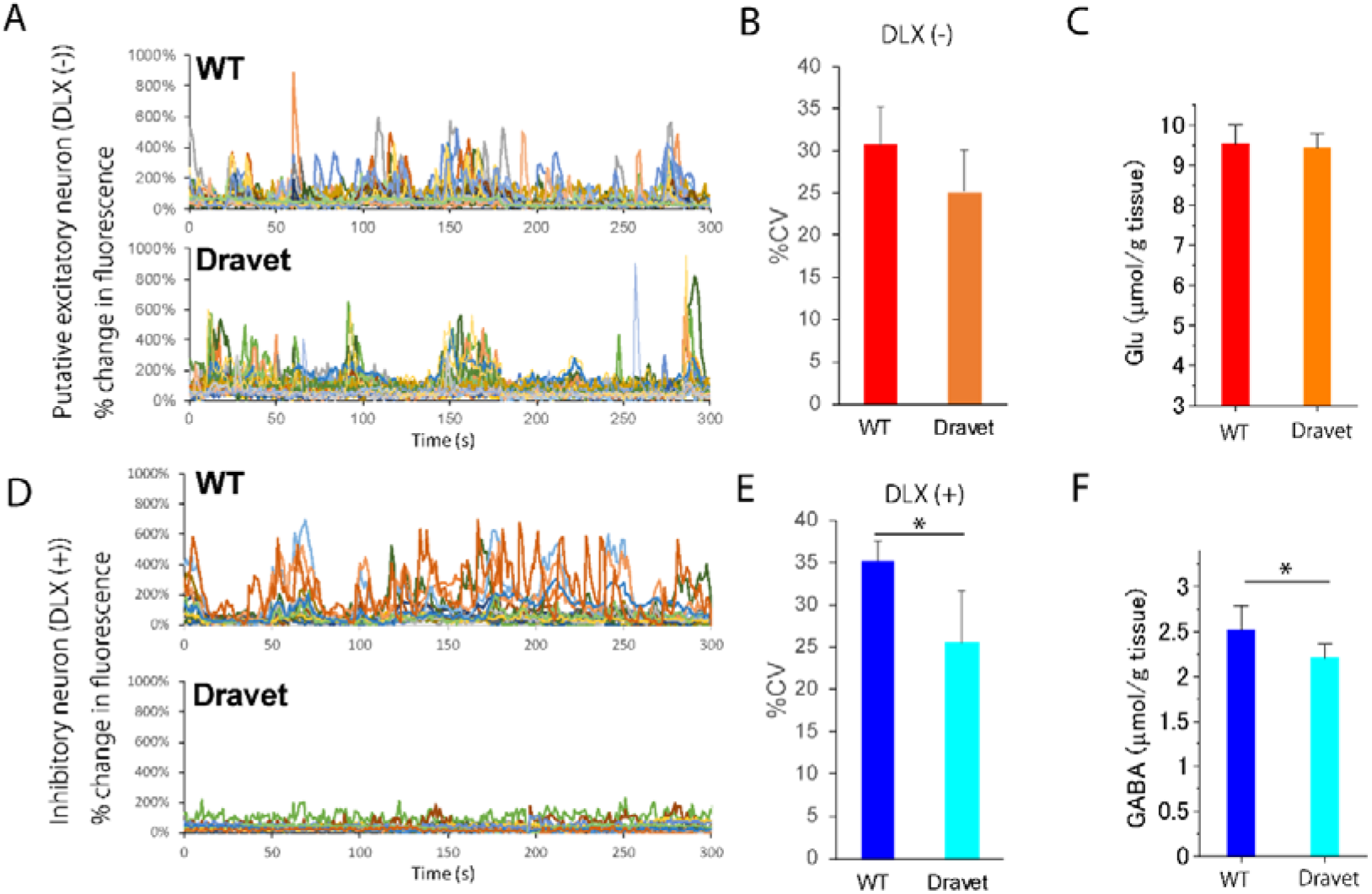
MRS-measured Glu and GABA and neural activities in a mouse model of Dravet syndrome with spontaneous brain activities. (A, B, C) Ca signals of putative excitatory neurons imaged by two-photon microscopy showed no significant difference between the two groups (paired t-test, n = 4 mice, t = −1.971, p = 0.143), which was in accordance with the unchanged MRS-measured Glu levels (two-sample t-test, n = 7, t = 0.0659, p = 0.9482; Fig. 4C). (D, E, F) Ca signals of inhibitory neurons decreased significantly in a mouse model of Dravet syndrome (two-sample t-test, n = 41-62 cells, t = −4.168, p=6.628e-5), which was in accordance with the results of MRS-measured GABA (two-sample t-test, n = 8, t = 3.0098, p = 0.008; (F)). In the Fig.4B and E, the percent coefficient of variance (%CV), which is defined as the ratio of standard deviation to the mean of fluorescence intensities, is shown. Error bars indicate standard deviation.

## Discussion

Despite the increasing number of findings based on MRS, one open question remains unresolved: do MRS-measured Glu and GABA in a brain region reflect activities of excitatory and inhibitory neurons, respectively, in the region? This uncertainty has been one of the main limitations to understanding the relationship between behavior and brain processing that underlies MRS signals. Using MRS and two-photon microscopy in the mouse brain, the present work collectively demonstrated that MRS-measured Glu and GABA levels can reflect excitatory and inhibitory neural activities, respectively, in the awake and behaving conditions.

### Awake MRS for exploring the neural basis of brain activities in human

An increasing number of human MR studies for exploring brain function using MRS have been performed ^1, 3, 30, 31^; however, no human studies demonstrated the underlying neural basis of altered neurochemicals measured by MRS. Thus, we utilized a custom-made awake mouse restraint device to perform whisker stimulation for causing neural activities, which was confirmed by an awake two-photon imaging system, in the barrel cortex of a mouse brain. Using those devices, we successfully detected neural activities as well as Glu and GABA level alterations, both of which were significantly suppressed by isoflurane anesthesia (Supplementary Fig.3). Collectively, this work using awake mouse MRS provides the rationale for human MRS studies, most of which are performed in the awake condition, for exploring the neurochemical basis of cognition and learning ^1, 5^, which has not been possible by other modalities.

### Spontaneous neural activities are associated with MRS-measured Glu and GABA levels

The results obtained from the Dravet mouse model in this work indicated that MRS-measured Glu and GABA at quiet rest reflect neural excitatory and inhibitory activities, respectively. This may also imply that awake MRS is sensitive enough to detect neuronal activities via Glu and GABA level alterations at quiet rest, using persistent neuronal activities during a resting condition. This notion is supported by previous works that demonstrated the relationship between Glu and GABA levels and altered signals of the resting state fMRI ^3, 32^. The other previous work, which detected different Glu levels according to the open/closed eye status ^31^, also suggested the sensitivity of the MRS measurement during the awake condition. This line of evidence indicates that awake MRS is capable of investigating brain function using Glu and GABA alterations even during quiet rest.

### Alterations of MRS-measured Glu and GABA levels in various conditions

MRS-measured Glu and GABA reflect the total Glu and GABA levels in the brain region, which are derived from all compartments, including cytosol, synaptic vesicles, and extracellular space in the VOI. It is assumable that there are different levels of contribution from each compartment depending on the brain conditions such as resting/active, and pathological/physiological conditions. In the case of Glu levels at quiet rest, MRS-measured Glu may be prone to reflect the synaptic density, given the abundant localization of Glu in presynaptic vesicles ^33^. Indeed, a previous report about Alzheimer’s disease model mice demonstrated that synaptic loss was associated with decreased MRS-measured Glu^34^. On the other hand, the behavior of Glu levels may be different in the diseased brain compared to the physiological condition, as was shown in fMRI studies of the diseased brain ^35^. This notion is supported by the fact that Glu can be consumed when there is not enough glucose as brain fuel ^36^. Nevertheless, awake MRS in the diseased brain, such as in Alzheimer’s disease model mice, would be intriguing, since excitation and inhibition balance (E/I balance) disruption exists in Alzheimer’s disease ^37^ as an important potential pathological mechanism.

In contrast to the complexity of Glu localization, a large portion of GABA in the brain under basal conditions is reported to localize in the cytoplasm ^38^, suggesting that MRS-measured GABA may reflect the GABA pool in the cytoplasm. While the functional significance of the GABA pool has remained unknown ^39^, several previous studies reported that baseline GABA levels in humans were associated with brain functions ^40-42^. The association between MRS-measured GABA and inhibitory neural activities at quiet rest in a Dravet mouse model ^43^ in the present work thus highlighted the importance of baseline GABA levels, which may be attributed to the cytoplasmic GABA and reflect inhibitory neural activities.

## Conclusion

Our work indicates that MRS-measured Glu and GABA could be an index of excitatory and inhibitory activities, respectively, in both spontaneous and task-evoked neural activities. By using MRS in an awake condition, we can monitor the E/I balance in various conditions, such as epilepsy and neuropsychiatric/neurological disorders, which provide us with important information in clinical studies. Also, using awake MRS in mice, we may address physiological and/or pathological questions by combining with other modalities such as positron emission tomography ^44-46^. Moreover, this method can be combined with optogenetics ^47^ and chemogenetics for further validating the mechanistic link between MRS findings and neural activities. Lastly, given that glial cells, such as astrocytes, are also known to contribute to Glu metabolism ^48^, there are likely other factors, particularly in the pathological conditions, that contribute to MRS neurotransmitter measures besides neuronal activity, which should be addressed in future study.

## Acknowledgements

We would like to express our appreciation to Ms Kanami Ebata, Mr Takeharu Minamihisamatsu, Mr Takahiro Shimizu and other colleagues who helped our experiments. We also appreciate the valuable comments about this manuscript by Dr. Takafumi Minamimoto. This project is supported by Kakenhi Grant Number 19K08005. Y Takado and KS are partially supported by Kakenhi Grant Number 19H01041. This research is also supported by AMED under Grant Number 17dm0107066h (IA) and partly supported by COI program (JPMJCE1305, IA) by JST.

## Author contribution statement

Y. Takado, HT and MH came up with the concept of the paper and designed experimental procedures. Y. Takado, H.T., K Sampei, T.U., S.U., N.N., S.S. performed MRS experiment. JN made a basis set for the MRS analysis. H.T., T.U., M.T., M.S., M.O., J.M., Y. Tomita designed and performed two-photon imaging experiments. K.N. and Y.O. designed a homemade laser system for mesoscopic imaging. Y. Takado and K. Shibata wrote the original draft of the paper. All authors gave comments on the manuscript and approved the final version of the paper.

## References

1. Shibata K, Sasaki Y, Bang JW, Walsh EG, Machizawa MG, Tamaki M et al. Overlearning hyperstabilizes a skill by rapidly making neurochemical processing inhibitory-dominant. Nat Neurosci 2017; 20(3): 470–475.

2. Kolasinski J, Logan JP, Hinson EL, Manners D, Divanbeighi Zand AP, Makin TR et al. A Mechanistic Link from GABA to Cortical Architecture and Perception. Current biology: CB 2017; 27(11): 1685-1691.e3.

3. Stagg CJ, Bachtiar V, Amadi U, Gudberg CA, Ilie AS, Sampaio-Baptista C et al. Local GABA concentration is related to network-level resting functional connectivity. eLife 2014; 3: e01465.

4. Stagg CJ, Bachtiar V, Johansen-Berg H. The role of GABA in human motor learning. Current biology: CB 2011; 21(6): 480–4.

5. Jocham G, Hunt LT, Near J, Behrens TE. A mechanism for value-guided choice based on the excitation-inhibition balance in prefrontal cortex. Nat Neurosci 2012; 15(7): 960–1.

6. Muthukumaraswamy SD, Edden RA, Jones DK, Swettenham JB, Singh KD. Resting GABA concentration predicts peak gamma frequency and fMRI amplitude in response to visual stimulation in humans. Proceedings of the National Academy of Sciences of the United States of America 2009; 106(20): 8356–61.

7. Gaetz W, Edgar JC, Wang DJ, Roberts TP. Relating MEG measured motor cortical oscillations to resting gamma-aminobutyric acid (GABA) concentration. NeuroImage 2011; 55(2): 616–21.

8. Takuwa H, Matsuura T, Obata T, Kawaguchi H, Kanno I, Ito H. Hemodynamic changes during somatosensory stimulation in awake and isoflurane-anesthetized mice measured by laser-Doppler flowmetry. Brain Res 2012; 1472: 107–12.

9. Makaryus R, Lee H, Yu M, Zhang S, Smith SD, Rebecchi M et al. The metabolomic profile during isoflurane anesthesia differs from propofol anesthesia in the live rodent brain. Journal of cerebral blood flow and metabolism: official journal of the International Society of Cerebral Blood Flow and Metabolism 2011; 31(6): 1432–42.

10. Stagg CJ, Bestmann S, Constantinescu AO, Moreno LM, Allman C, Mekle R et al. Relationship between physiological measures of excitability and levels of glutamate and GABA in the human motor cortex. The Journal of physiology 2011; 589(Pt 23): 5845–55.

11. Kolasinski J, Hinson EL, Divanbeighi Zand AP, Rizov A, Emir UE, Stagg CJ. The dynamics of cortical GABA in human motor learning. The Journal of physiology 2019; 597(1): 271–282.

12. Öz G, Alger JR, Barker PB, Bartha R, Bizzi A, Boesch C et al. Clinical Proton MR Spectroscopy in Central Nervous System Disorders. Radiology 2014; 270(3): 658–679.

13. Takado Y, Igarashi H, Terajima K, Shimohata T, Ozawa T, Okamoto K et al. Brainstem metabolites in multiple system atrophy of cerebellar type: 3.0-T magnetic resonance spectroscopy study. Movement Disorders 2011; 26(7): 1297–1302.

14. Takado Y, Terajima K, Ohkubo M, Okamoto K, Shimohata T, Nishizawa M et al. Diffuse brain abnormalities in myotonic dystrophy type 1 detected by 3.0 T proton magnetic resonance spectroscopy. European neurology 2015; 73(3-4): 247–56.

15. Sofroniew NJ, Flickinger D, King J, Svoboda K. A large field of view two-photon mesoscope with subcellular resolution for in vivo imaging. eLife 2016; 5.

16. Floyer-Lea A, Wylezinska M, Kincses T, Matthews PM. Rapid modulation of GABA concentration in human sensorimotor cortex during motor learning. Journal of neurophysiology 2006; 95(3): 1639–44.

17. Yu FH, Mantegazza M, Westenbroek RE, Robbins CA, Kalume F, Burton KA et al. Reduced sodium current in GABAergic interneurons in a mouse model of severe myoclonic epilepsy in infancy. Nature neuroscience 2006; 9(9): 1142–9.

18. Matsushima Y. A new mouse model for severe myoclonic epilepsy in infancy: positional cloning of the causative gene and pathological analysis. Ann.Rep.Jpn.Epi.Res.Found. 2015; 26: 69–76.

19. Takuwa H, Tajima Y, Kokuryo D, Matsuura T, Kawaguchi H, Masamoto K et al. Hemodynamic changes during neural deactivation in awake mice: a measurement by laser-Doppler flowmetry in crossed cerebellar diaschisis. Brain Res 2013; 1537: 350–5.

20. Chen TW, Wardill TJ, Sun Y, Pulver SR, Renninger SL, Baohan A et al. Ultrasensitive fluorescent proteins for imaging neuronal activity. Nature 2013; 499(7458): 295–300.

21. Dimidschstein J, Chen Q, Tremblay R, Rogers SL, Saldi GA, Guo L et al. A viral strategy for targeting and manipulating interneurons across vertebrate species. Nature neuroscience 2016; 19(12): 1743–1749.

22. McClure C, Cole KL, Wulff P, Klugmann M, Murray AJ. Production and titering of recombinant adeno-associated viral vectors. Journal of visualized experiments: JoVE 2011; (57): e3348.

23. Tomita Y, Kubis N, Calando Y, Tran Dinh A, Meric P, Seylaz J et al. Long-term in vivo investigation of mouse cerebral microcirculation by fluorescence confocal microscopy in the area of focal ischemia. J Cereb Blood Flow Metab 2005; 25(7): 858–67.

24. Takuwa H, Masamoto K, Yamazaki K, Kawaguchi H, Ikoma Y, Tajima Y et al. Long-term adaptation of cerebral hemodynamic response to somatosensory stimulation during chronic hypoxia in awake mice. J Cereb Blood Flow Metab 2013; 33(5): 774–9.

25. Zaouter Y, Papadopoulos DN, Hanna M, Boullet J, Huang L, Aguergaray C et al. Stretcher-free high energy nonlinear amplification of femtosecond pulses in rod-type fibers. Optics letters 2008; 33(2): 107–9.

26. Nagashima K, Ochi Y, Itakura R. Statistical effects of optical parametric noise on signal pulses in a synchronously pumped optical parametric oscillator. J. Opt. Soc. Am. B 2019; 36(12): 3389–3394.

27. Nagashima K, Ochi Y, Itakura R. Optical parametric oscillator pumped by a 100-kHz burst-mode Yb-doped fiber laser. Optics letters 2020; 45(3): 674–677.

28. Ogawa S, Lee TM, Kay AR, Tank DW. Brain magnetic resonance imaging with contrast dependent on blood oxygenation. Proc Natl Acad Sci U S A 1990; 87(24): 9868–72.

29. Provencher SW. Automatic quantitation of localized in vivo1H spectra with LCModel. NMR in Biomedicine 2001; 14(4): 260–264.

30. Kurcyus K, Annac E, Hanning NM, Harris AD, Oeltzschner G, Edden R et al. Opposite Dynamics of GABA and Glutamate Levels in the Occipital Cortex during Visual Processing. The Journal of neuroscience: the official journal of the Society for Neuroscience 2018; 38(46): 9967–9976.

31. Lynn J, Woodcock EA, Anand C, Khatib D, Stanley JA. Differences in steady-state glutamate levels and variability between ïnon-task-activeï conditions: Evidence from (1)H fMRS of the prefrontal cortex. Neuroimage 2018; 172: 554–561.

32. Levar N, Van Doesum TJ, Denys D, Van Wingen GA. Anterior cingulate GABA and glutamate concentrations are associated with resting-state network connectivity. Scientific reports 2019; 9(1): 2116.

33. Nedergaard M, Takano T, Hansen AJ. Beyond the role of glutamate as a neurotransmitter. Nature reviews. Neuroscience 2002; 3(9): 748–55.

34. Crescenzi R, DeBrosse C, Nanga RP, Reddy S, Haris M, Hariharan H et al. In vivo measurement of glutamate loss is associated with synapse loss in a mouse model of tauopathy. NeuroImage 2014; 101: 185–92.

35. Schridde U, Khubchandani M, Motelow JE, Sanganahalli BG, Hyder F, Blumenfeld H. Negative BOLD with large increases in neuronal activity. Cerebral cortex (New York, N.Y.: 1991) 2008; 18(8): 1814–27.

36. Behar KL, den Hollander JA, Petroff OA, Hetherington HP, Prichard JW, Shulman RG. Effect of hypoglycemic encephalopathy upon amino acids, high-energy phosphates, and pHi in the rat brain in vivo: detection by sequential 1H and 31P NMR spectroscopy. Journal of neurochemistry 1985; 44(4): 1045–55.

37. Ren SQ, Yao W, Yan JZ, Jin C, Yin JJ, Yuan J et al. Amyloid beta causes excitation/inhibition imbalance through dopamine receptor 1-dependent disruption of fast-spiking GABAergic input in anterior cingulate cortex. Scientific reports 2018; 8(1): 302.

38. Asada H, Kawamura Y, Maruyama K, Kume H, Ding RG, Kanbara N et al. Cleft palate and decreased brain gamma-aminobutyric acid in mice lacking the 67-kDa isoform of glutamic acid decarboxylase. Proceedings of the National Academy of Sciences of the United States of America 1997; 94(12): 6496–9.

39. Maddock RJ, Buonocore MH. MR spectroscopic studies of the brain in psychiatric disorders. Current topics in behavioral neurosciences 2012; 11: 199–251.

40. Boy F, Evans CJ, Edden RA, Singh KD, Husain M, Sumner P. Individual differences in subconscious motor control predicted by GABA concentration in SMA. Current biology: CB 2010; 20(19): 1779–85.

41. Puts NA, Edden RA, Evans CJ, McGlone F, McGonigle DJ. Regionally specific human GABA concentration correlates with tactile discrimination thresholds. The Journal of neuroscience: the official journal of the Society for Neuroscience 2011; 31(46): 16556–60.

42. Sumner P, Edden RA, Bompas A, Evans CJ, Singh KD. More GABA, less distraction: a neurochemical predictor of motor decision speed. Nature neuroscience 2010; 13(7): 825–7.

43. Richards KL, Milligan CJ, Richardson RJ, Jancovski N, Grunnet M, Jacobson LH et al. Selective NaV1.1 activation rescues Dravet syndrome mice from seizures and premature death. Proceedings of the National Academy of Sciences of the United States of America 2018; 115(34): E8077–e8085.

44. Jauhar S, McCutcheon R, Borgan F, Veronese M, Nour M, Pepper F et al. The relationship between cortical glutamate and striatal dopamine in first-episode psychosis: a cross-sectional multimodal PET and magnetic resonance spectroscopy imaging study. The lancet. Psychiatry 2018; 5(10): 816–823.

45. Maruyama M, Shimada H, Suhara T, Shinotoh H, Ji B, Maeda J et al. Imaging of tau pathology in a tauopathy mouse model and in Alzheimer patients compared to normal controls. Neuron 2013; 79(6): 1094–108.

46. Ishikawa A, Tokunaga M, Maeda J, Minamihisamatsu T, Shimojo M, Takuwa H et al. In Vivo Visualization of Tau Accumulation, Microglial Activation, and Brain Atrophy in a Mouse Model of Tauopathy rTg4510. Journal of Alzheimer’s disease: JAD 2018; 61(3): 1037–1052.

47. Just N, Faber C. Probing activation-induced neurochemical changes using optogenetics combined with functional magnetic resonance spectroscopy: a feasibility study in the rat primary somatosensory cortex. Journal of neurochemistry 2019; 150(4): 402–419.

48. Danbolt NC. Glutamate uptake. Progress in neurobiology 2001; 65(1): 1–105.

