## Supplemental Figures for "MRS-measured Glutamate versus GABA reflects excitatory versus inhibitory neural activities in awake mice"

**Supplementary Figures**

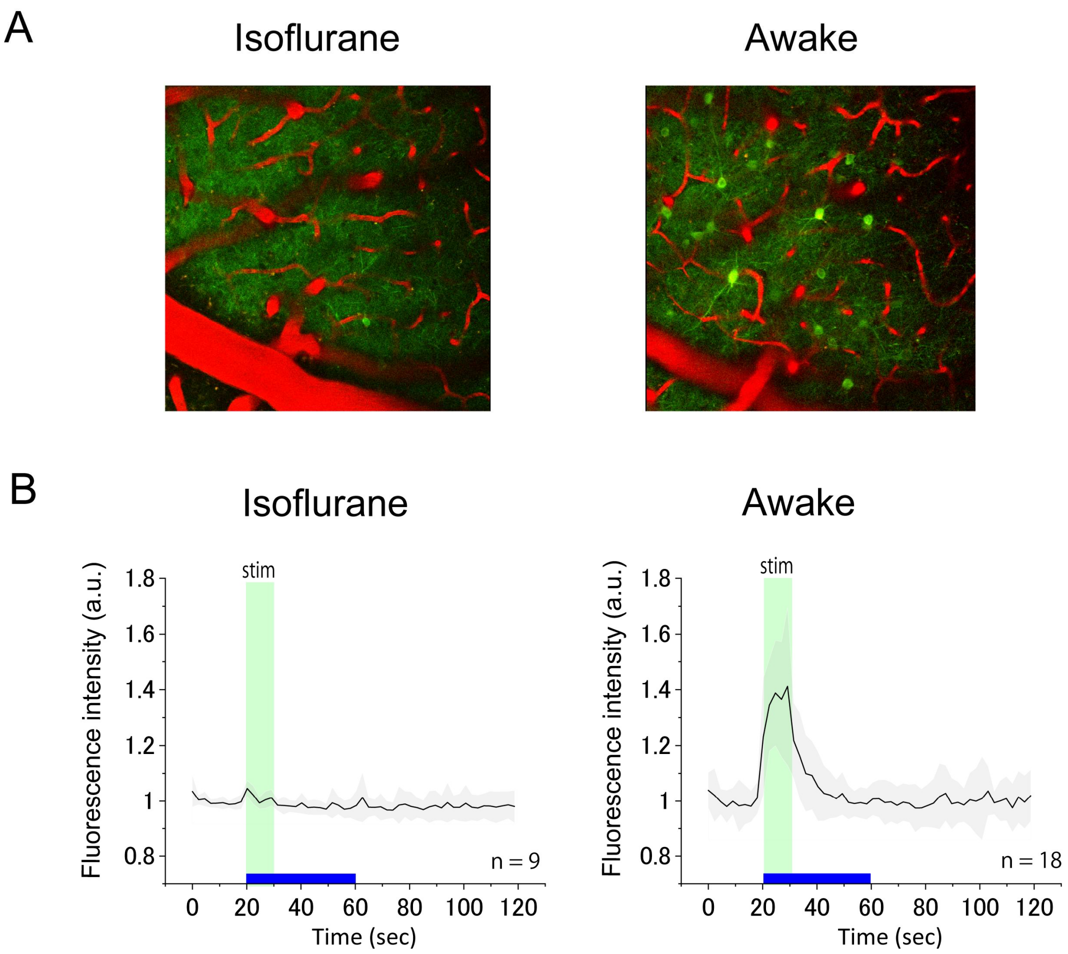


**Supplementary Fig.1. Comparison of neural activity between anesthetic and awake conditions**

Whisker stimulation on mice was performed for 10 sec by air-puff at 1 Hz. While no significant alterations of Ca signal intensities were found under anesthetic condition (A, B left), alterations of Ca signal intensities during the awake condition were evident (A, B right). Each black line shows the average fluorescence intensities of multiple neurons (n = 8 with isoflurane, n = 18 for awake condition). A gray area indicates standard deviation.

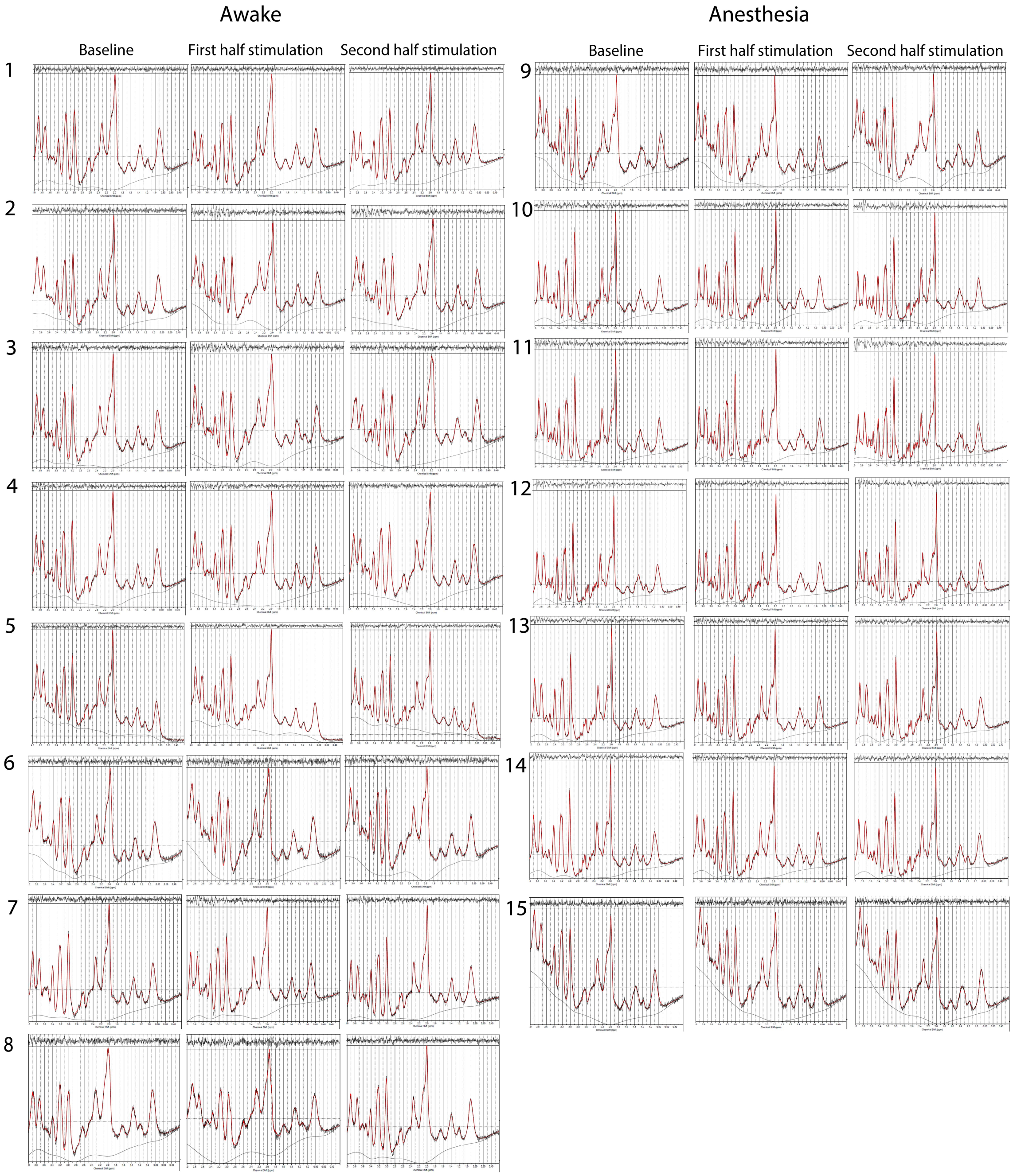


**Supplementary Fig.2. LCModel fitted spectra of all MRS experiments**

All LCModel fitted spectra are shown. The left group (1 to 8) shows spectra in the awake condition. The right group (9 to 15) shows spectra in the anesthetic condition.

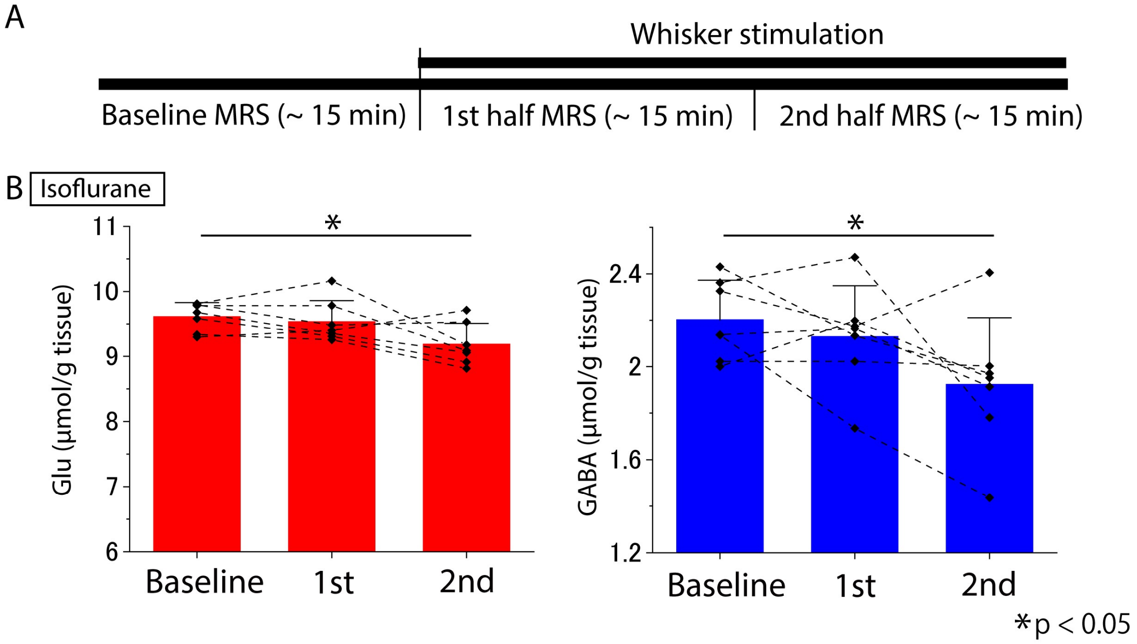


**Supplementary Fig.3 Alterations of Glu and GABA levels during whisker stimulation measured by MRS with anesthesia.** (A) The MRS protocol is shown. (B) Quantitative alterations of Glu and GABA levels at the anesthetic condition are shown. The Glu and GABA levels were significantly decreased compared to baseline in the second half session (Glu; one-way repeated-measures ANOVA with Bonferroni post-hoc test (rm-ANOVA), $n = 7$, $t = 3.055$, $p = 0.030$, GABA; rm-ANOVA, $n = 7$, $t = 2.737$, $p = 0.018$). Error bars indicate standard deviation.

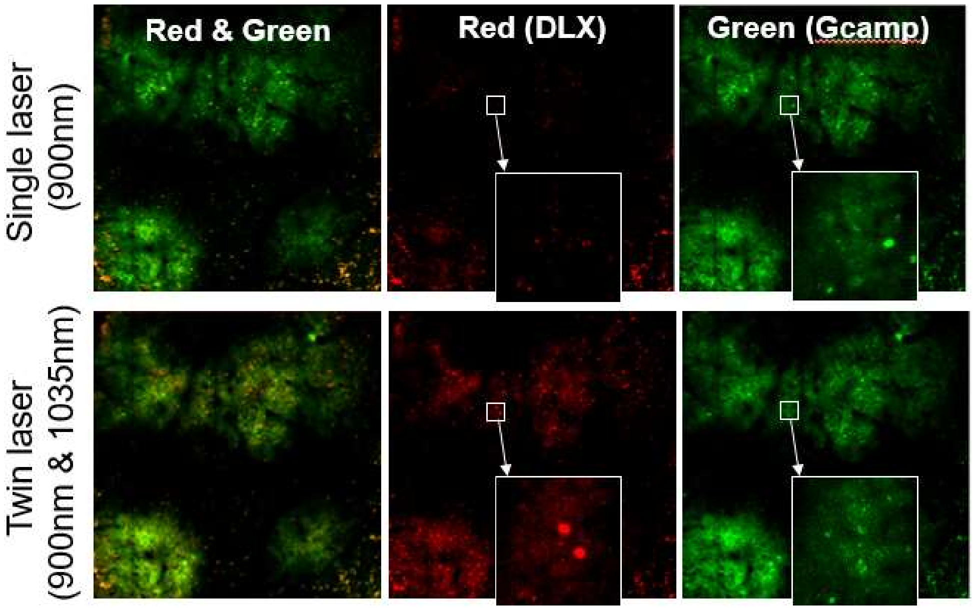


**Supplementary Fig.4**

Upper and lower figures show two-photon images with a single laser device and a twin laser device, respectively. Regarding the twin laser system (lower figures), while the added custom-made laser oscillator increased intensities of DLX fluorescence, the other commercial laser oscillator increased the intensities of Gcamp brightness. As a result, the twin laser system can discriminate better between Gcamp and DLX fluorescence intensities, resulting in more evident visualization of DLX positive neurons.
